# Cas9-mediated gene-editing in the malaria mosquito *Anopheles stephensi* by ReMOT Control

**DOI:** 10.1101/775312

**Authors:** Vanessa M. Macias, Sage McKeand, Duverney Chaverra-Rodriguez, Grant L. Hughes, Aniko Fazekas, Sujit Pujhari, Nijole Jasinskiene, Anthony A. James, Jason L. Rasgon

## Abstract

Innovative tool development is essential for continued advancement in malaria control and depends on a deeper understanding of the molecular mechanisms that govern transmission of malaria parasites by *Anopheles* mosquitoes. Targeted disruption of genes in mosquito vectors is a powerful method to uncover the underlying biology of vector-pathogen interactions, and genome manipulation technologies can themselves form the basis of mosquito and pathogen control strategies. However, the embryo injection methods used to genetically manipulate mosquitoes, and in particular *Anopheles* species, are difficult and inefficient, particularly for non-specialist laboratories. We have adapted a strategy called ReMOT Control (<u>Re</u>ceptor-<u>m</u>ediated <u>O</u>vary <u>T</u>ransduction of <u>C</u>argo) to deliver the Cas9 ribonucleoprotein complex to adult mosquito ovaries and generate targeted and heritable mutations in the malaria vector *Anopheles stephensi*. We found that gene editing by ReMOT Control in *Anopheles* mosquitoes was comparable to the technique in *Ae. aegypti* and as efficient in editing as standard embryo injections. The adaptation of this technology to *Anopheles* mosquitoes opens up the power of reverse genetics to malaria vector labs that do not have the equipment or technical expertise to perform embryo injections and establishes the flexibility of ReMOT Control for gene-editing in non-*Aedes* species.

## Introduction

To solve the growing problem of mosquito-borne disease, we need improved and more efficient gene editing methods. CRISPR/Cas9-based targeted disruption of genes in mosquito vectors of human pathogens is a powerful method to uncover underlying aspects of mosquito pathogen transmission biology that can be targeted for basic biological understanding and for mosquito control efforts^1–5^. Genome manipulation technologies can themselves form the basis of mosquito and pathogen control strategies^6,7^. While many techniques based on CRISPR/Cas9 have improved studies in mosquito genetics, these techniques rely on embryo injection of the Cas9 ribonucleoproten complex (RNP), which requires specialized and expensive equipment and training. To get around this bottleneck, a strategy termed ReMOT Control (Receptor-mediated Ovary Transduction of Cargo) was developed (8) for delivery of gene-editing moieties to the arthropod germline from the hemocoel, allowing targeted and heritable mutations to be made by adult injection instead of embryo microinjection. Proof-of-principle experiments validating ReMOT Control were conducted in *Aedes aegypti*, but extension of the method to species more recalcitrant to editing (such as *Anopheles* mosquitoes) was not established^8^.

In this study, we have adapted the ReMOT Control technique for efficient gene editing in the human malaria vector *Anopheles stephensi*. We found that once optimized, gene editing by ReMOT Control in *Anopheles* mosquitoes was comparable to the technique in *Ae. aegypti* and as efficient in editing as standard embryo injections. As Anophelines are substantially more difficult to manipulate at the embryo stage than *Aedes* mosquitoes, ReMOT Control represents a much-needed improvement in gene editing for malaria vectors. We expect that ReMOT Control gene editing will become as commonly used for reverse genetics in *Anopheles* mosquitoes as it has become in Aedine vectors of human arboviruses.

## Results and Discussion

### *ECFP* targeting, mutation detection scheme, and validation by embryo microinjection

To establish a method for site-specific editing in *An. stephensi*, we used ReMOT Control to target one of the two fluorescent marker genes in the transgenic *An. stephensi* line VgCp26.10^7^, which offered several advantages for establishment of mutagenesis methods over the endogenous eye color genes previously used. The gene encoding *kynurenine 5-monooxygenase* (*kmo* or *kynurenine hydroxylase, kh*) was used as a visible marker to develop both embryo injection and adult injection methods in *Ae. aegypti*^8,9^.

However, the *kmo* knockout is associated with substantially decreased fitness in *An. stephensi*, while the VgCp26.10 transgenic line is robust and is not likely to suffer from the phenotypic loss of a fluorescent marker^7,10^. A mutation in the one dominant visible fluorescent marker gene (*enhanced cyan fluorescence protein, ECFP*) is easy to detect in hemizygous live mosquitoes by fluorescence imaging while simultaneous expression of the intact linked marker gene (*Discosoma species Red, DsRed*) confirms presence of the transgene rather than the wild-type empty locus^11^.

Initial validation of targeted knockdown of *ECFP* in the VgCp26.10 transgenic line by embryo microinjection provided a benchmark to assess ReMOT Control knockout efficiency. Homozygous transgenic embryos (generation 0, G_0_) were injected with Cas9 plasmid expression cassette, mRNA, or protein, and three single guide RNAs originally designed to target EGFP in human cells^12^ but which also cut ECFP due to a high degree of sequence conservation between the two genes. Surviving G_0_ embryos injected with 200 ng/µL Cas9 protein and 300 ng/µL sgRNAs produced G_1_ offspring lacking cyan fluorescence in the eyes but expressing DsRed in four out of 32 single male G_1_ families and one female pool, resulting in frequencies of 12.5% families producing edited offspring and 1.5-24% of G_1_ offspring edited within those families (Table 1 and Supplemental Table 1). *ECFP* knockout individuals comprised 1.5% of total G_1_ screened across all families. Two male families produced individuals with loss of both cyan and red fluorescence, indicating that a modification occurred that not only interrupted *ECFP* expression, but also interrupted expression of the *DsRed* marker that was over 7 kilobases away (Figure 1). Such large deletions are not unprecedented, but are likely under-reported in studies where PCR amplicons of limited size are used to detect deletions^13^. Injection of embryos with a reduced concentration of total gRNA (75ng/µL) produced fewer visually detectible mutations; G_1_ individuals with *ECFP* disruption were only identified from one pool of females out of 11 total female pools and single male families and constituted 0.13% of G_1_ individuals screened (Supplemental Tables 1 and 2). No knock-outs were detectable in G_1_ larvae when G_0_ embryos were injected with 500 ng/µL of Cas9 transcript or 500 ng/µL of plasmid with Cas9 expressed from the Hsp70 promoter (pDCC6,^14^) (Supplemental Table 3).

**Table 1:** Injection data and comparison of ECFP mutation by adult and embryo injections.

| Injection Mix |  |  | G<sub>1</sub> |  |  |  | G<sub>0</sub> Fluorescence Phenotype |  |  |  |  | G<sub>1</sub> Fluorescence Phenotype |  |  |  |  |
| --- | --- | --- | --- | --- | --- | --- | --- | --- | --- | --- | --- | --- | --- | --- | --- | --- |
| Protein | 3 sgCFP | Saponin | G<sub>1</sub> Cross | No. Inj | Inj Time PBM | No. on EC day | No. | Cyan+<br>DsRed+ | Cyan-<br>DsRed- | Cyan-<br>DsRed+ | Cyan Mosaic<br>DsRed+ | G<sub>0</sub> C+R+<br>Outcross | No. | Cyan+<br>DsRed+ | Cyan-<br>DsRed- | Cyan-<br>DsRed+ |
| Adult injections: P2C-Cas9/gRNA mix injections into VgCp26.10 colony females (G<sub>1</sub>) crossed with wild type males |  |  |  |  |  |  |  |  |  |  |  |  |  |  |  |  |
| 0.5 µg/µL | 0.75 µg/µL | + | VgCp26.10 Females X WT Males | 300 | 24hrs | 212 | 425 | 106 | 317 | 2<br>(0.47%) | 0 | Female | 295 | 103 | 192 | 0 |
|  |  |  |  |  |  |  |  |  |  |  |  | Male | 622 | 324 | 298 | 0 |
| 0.75 µg/µL | 1.2 µg/µL | + | VgCp26.10 Females X WT Males | 83 | 48 hrs | 60 | 194 | 160 | 24 | 10<br>(5.2%) | 1*<br>(1.2%) | Female | 52 | 26 | 26 | 0 |
|  |  |  |  |  |  |  |  |  |  |  |  | Male | 102 | 38 | 53 | 11 (10.8%) |
| 0.75 µg/µL | 1.2 µg/µL | + | VgCp26.10 Females X WT Males | 63 | 47 hrs | 33 | 161 | 52 | 107 | 2<br>(1.2%) | 1**<br>(0.62%) | Female | 61 | 29 | 32 | 0 |
|  |  |  |  |  |  |  |  |  |  |  |  | Male | 84 | 35 | 49 | 0 |
| 0.75 µg/µL | 1.2 µg/µL | + | VgCp26.10 Females X WT Males | 78 | 46 hrs | 46 | 71 | 41 | 30 | 0 | 2**<br>(2.8%) | Female | 113 | 30 | 83 | 0 |
|  |  |  |  |  |  |  |  |  |  |  |  | Male |  |  | ND |  |
| Sap+ Totals: |  |  |  | 524 |  | 351 | 851 | 359 | 478 | 14 (1.7%) | 4 (0.47%) |  | 1329 | 585 | 733 | 11 (0.82%) |
| 0.75 µg/µL | 1.1 µg/µL | - | VgCp26.10 Females X WT Males | 97 | 28 hrs | 78 | 184 | 114 | 70 | 0 | 0 | Female | 217 | 96 | 118 | 3 (1.4%) |
|  |  |  |  |  |  |  |  |  |  |  |  | Male | 97 | 49 | 48 | 0 |
| 0.75 µg/µL | 1.1 µg/µL | - | VgCp26.10 Females X WT Males | 71 | 25 hrs | 43 | 173 | 61 | 112 | 0 | 0 | Female | 41 | 15 | 26 | 0 |
|  |  |  |  |  |  |  |  |  |  |  |  | Male | 207 | 109 | 98 | 0 |
| Sap - Totals: |  |  |  | 168 |  | 121 | 357 | 175 | 182 | 0 | 0 |  | 562 | 269 | 290 | 3 (0.53%) |
| Totals: |  |  |  | 692 |  | 472 | 1208 | 534 | 660 | 0 | 0 |  | 1891 | 854 | 1023 | 14 (0.74%) |
| Embryo injections: Cas9/gRNA mix injections into homozygous VgCp26.10 transgenic embryos (G<sub>0</sub>) |  |  |  |  |  |  |  |  |  |  |  |  |  |  |  |  |
| 0.2 µg/µL | 0.075 µg/µL | - | VgCp26.10 Intercross |  |  |  | 309 |  |  | Not Screened |  |  | 2320 | 2317 | 3 (0.13%) | 0 |
| 0.2 µg/µL | 0.3 µg/µL | - | VgCp26.10 Intercross |  |  |  | 382 |  |  | Not Screened |  |  | 5646 | 5506 | 52 (0.92%) | 88 (1.5%) |
| Totals: |  |  |  |  |  |  | 691 |  |  |  |  |  | 7966 | 7823 | 55 (0.69%) | 88 (1.1%) |

\*Individual had one cyan+ eye and one cyan- eye
\*\*these individuals had markedly reduced cyan fluorescence

**Figure 1:**
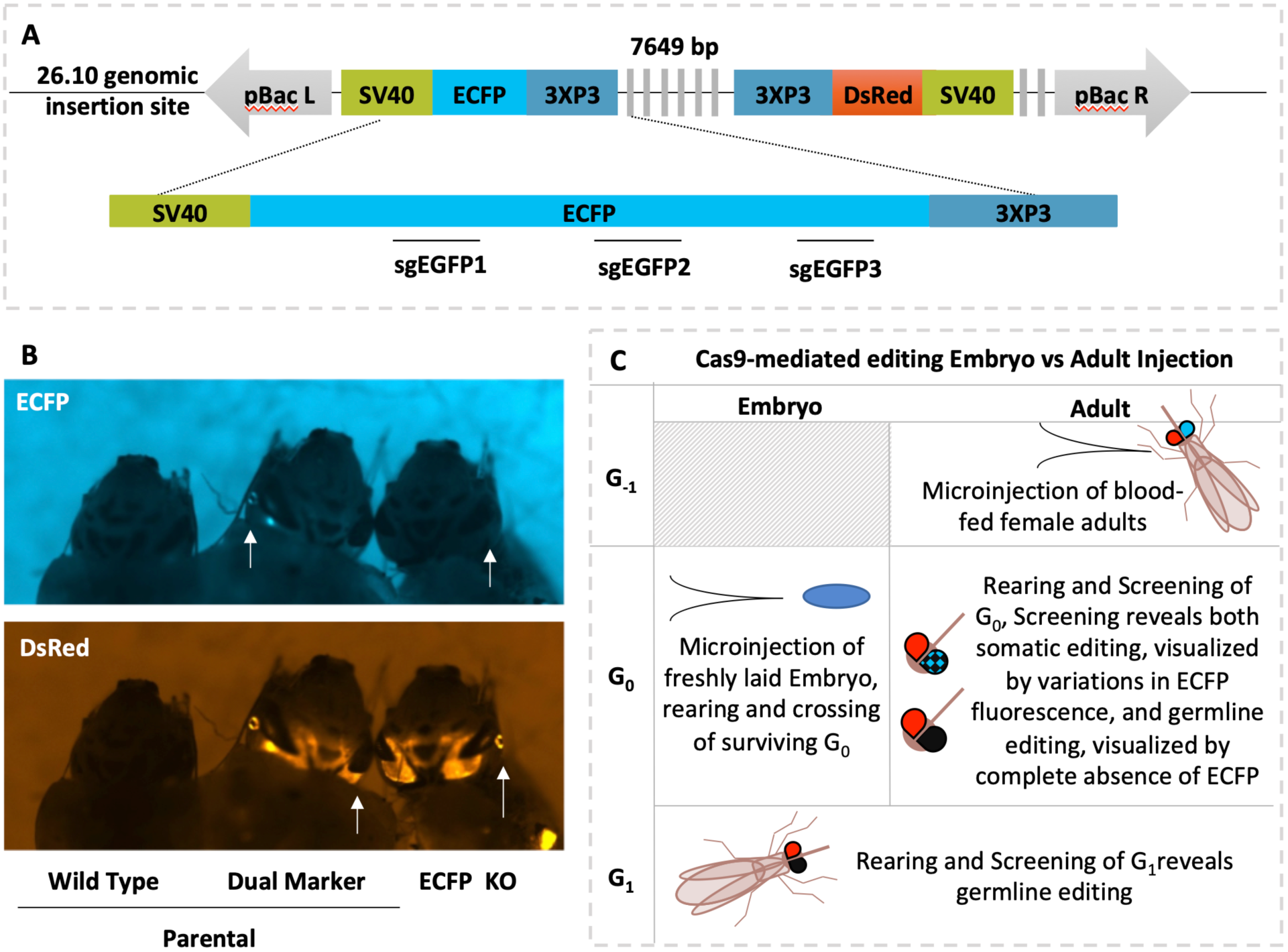
Schematic of knock-down approach. A) Schematic representation of transgene present in transgenic line VgCp26.10. The transgene has an ECFP and DsRed marker genes. Three guide RNAs (sgGFP) target Cas9 to the ECFP gene. B) Representative fluorescence image of larvae showing parental wild-type, parental transgenic and exceptional Cyan-, DsRed+ phenotypes. C) Schematic comparison of embryo and adult injection approaches for ECFP knock-out and detection of mutants The shaded box represents not done. Abbreviations are bp: base pair(s), pBac L/R: piggyBac left/right arm, SV40: simian virus 40 3’ untranslated region, DsRed: Discosoma species Red, ECFP: enhanced cyan fluorescent protein, sgGFP: single guide RNAs against GFP, KO: knock-out, G_x_: generation X.

### Optimization of injection components for adult injection

In order to accomplish heritable mutagenesis in *An. stephensi* by ReMOT Control, we utilized the ovary targeting P2C peptide derived from *Drosophila melanogaster* yolk protein 1 (DmYP1), which we recently reported in *Ae. aegypti* to mediate delivery of Cas9 RNP into vitellogenic oocytes^8^. However, we found that the *Aedes* injection parameters were not directly translatable to *An. stephensi*. Timing of injection of the P2C-Cas9 RNP relative to a blood-meal was reported to impact the developmental stage at which genome modifications are made in *Ae. aegypti.* Additionally *An. stephensi* have lower tolerance in general to injected components. We identified the injection conditions and timing that would maximize RNP uptake into the ovaries in a way that balanced survival and egg laying with concentrations of RNP effective for editing. We first injected females with P2C-EGFP fusion protein and visualized uptake into the ovary by fluorescence microscopy (Fig 2). Green fluorescence was more intense and present in all oocytes dissected from injected females when P2C-EGFP was injected within two days of blood feeding compared to decreased fluorescence when the protein was injected prior to a blood meal. Only 50% of ovary pairs had visible fluorescence in females injected 48 hours after a blood meal, perhaps because injected females were past the peak of vitellogenesis^15,16^ but those with visible fluorescence were comparable in intensity to earlier injections. Thus, experimental injections were performed within the first two days of a blood meal.

**Figure 2:**
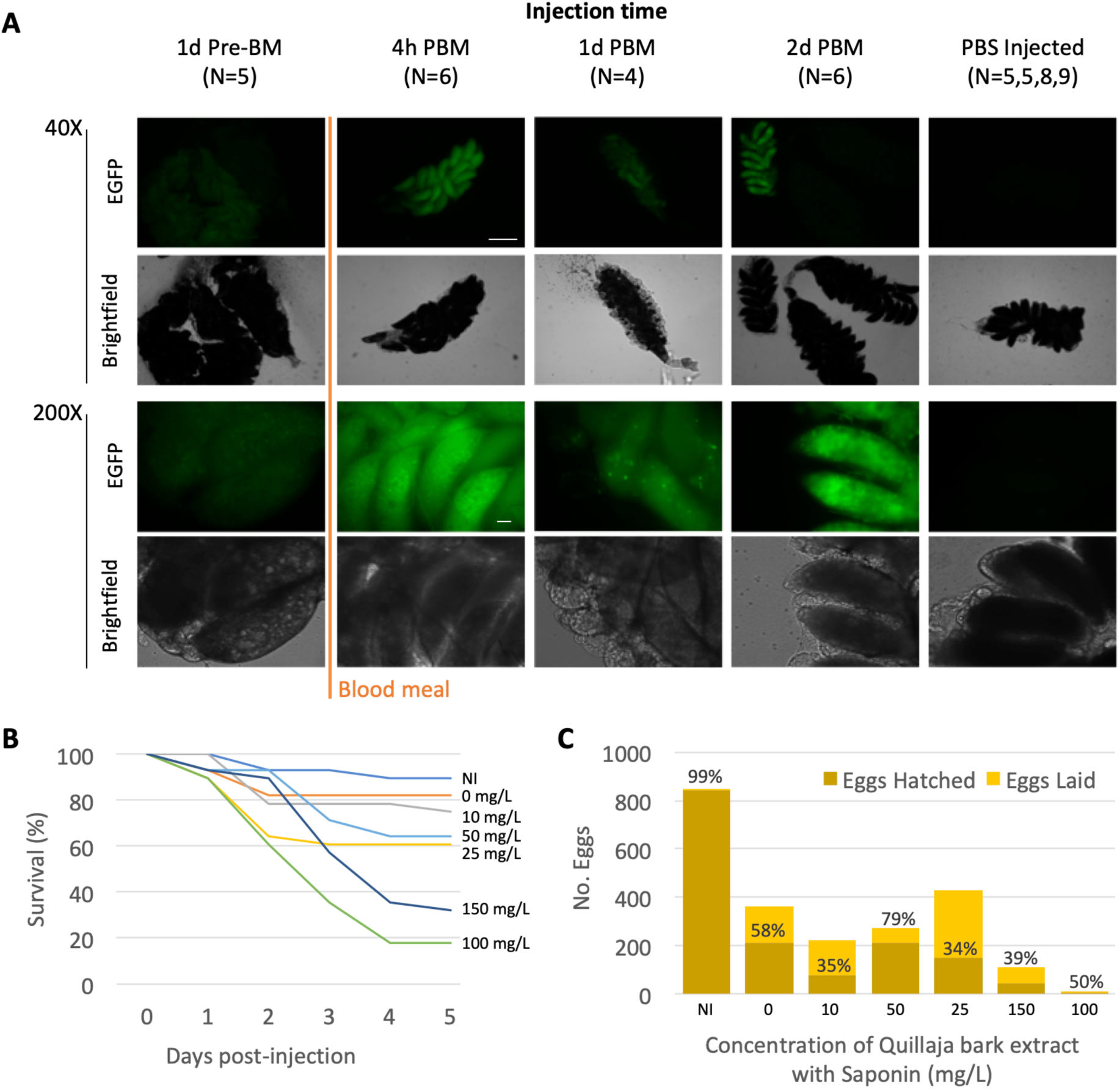
Optimization of injection components. A) Fluorescent and bright-field imaging of 72 hours (h) post-blood meal (PBM) ovaries following injection of P2C-EGFP at 1 day (d) Abbreviations are the same as Figure 1 and h: hour, N: number ovary pairs, pre-BM: before blood meal, PBS: phosphate buffered saline B) and C) Measurements of survival, egg laying and hatching following injections with varied concentrations of saponin.

In previous work, we used chloroquine to facilitate exit of the Cas9 RNP from the endosome after uptake, but we noted that saponin (*Quallaja* bark extract) also could be used for this purpose (8). For *Anopheles* ReMOT editing, we therefore investigated the use of saponin for endosome escape. In order to identify the highest concentration of saponin that would result in survival and viable offspring of injected adults, we performed a set of preliminary injections using different concentrations of the extract in 1X phosphate-buffered saline (PBS, Fig 2). Injection itself caused substantial decrease in survival and egg laying, but a concentration of 50 mg/L saponin could be injected and still have a high enough oviposition and hatch rate to have enough G_0_ progeny to screen (Figure 2). Importantly, we also determined that *An. stephensi* adults could not tolerate the dialysis buffer, which is used in the last step of purification of P2C-Cas9 from *E. coli* (and which was the buffer used for injections in *Ae. aegypti*). P2C-Cas9 precipitated out of solution when diluted in water or 1X PBS. We found that the buffer of P2C-Cas9 in complex with sgRNA did not precipitate and so could be column exchanged with water resulting in a final neutral injection solution with less than 5uM KCl remaining from the dialysis buffer, and that was tolerated by injected adults.

### *ECFP* marker knockout by ReMOT Control

P2C-Cas9 RNP with EGFP guide RNAs sgEGFP1, 2 and 3 were injected along with saponin (50 mg/L) into VgCp26.10 adult females (G_-1_) crossed to wild-type males, such that G_0_ and G_1_ progeny with parental phenotypes are either positive for both ECFP and DsRed, or wild-type. Expected mutant phenotypes from successful editing would be ECFP-, DsRed+. We recovered ECFP-, DsRed+ G_0_ individuals from three out of the four injection groups in which saponin was included, with a total of twelve ECFP-, DsRed + individuals out of 851 total G_0_ screened (1.4%). No mutant individuals were recovered from the G_0_ progeny in injections without saponin. The VgCp26.10 colony was not completely homozygous at the generation of adult injection, resulting in production of some wild-type G_0_, so the efficiency of editing is more accurately understood in terms of number of alleles edited rather than the number of individuals. In this case a total of 377 transgenic alleles were available for editing in the G_0_ progeny of saponin/P2C-Cas9 RNP injected G_-1_ females as visually detected by DsRed fluorescence in heterozygous G_1_ individuals and 14 (3.7%) were edited, whereas 0% of the 534 alleles were edited in the G_0_ progeny of G_-1_ females injected without saponin. This adjustment is relevant to the extension of this technique to the mutation of endogenous mosquito genes that will be present in two copies. In these applications, especially for genes that lack a visible phenotype, editing is sufficiently efficient to be able to screen for these mutants by PCR-based methods.

ECFP-, DsRed+ individuals that survived to adulthood were individually outcrossed to wild-type mosquitoes resulting in G_1_ progeny, of which approximately half were ECFP-,DsRed+ and half were wild-type, consistent with Mendelian segregation of two alleles (X^2^=0.0397, P=0.84) along with one transgenic mutant and one wild-type. We conclude from this that *ECFP* knockouts detected at G_0_ are complete knockouts and likely occurred as Cas9 cleavage of the DNA in the oocyte prior to embryogenesis (Table 2). Four G_0_ progeny were designated mosaics. One of these was ECFP positive in only one eye and the three others had markedly decreased ECFP fluorescence, which is unexpected in individuals with a transgene at a fixed location (Table 1). These individuals were outcrossed but produced no progeny.

**Table 2:** Outcross of cyan-, DsRed+ G_0_ demonstrates ECFP mutations are heritable

| G<sub>0</sub> Cross | G<sub>1</sub> Fluorescence Phenotype |  |  |  |
| --- | --- | --- | --- | --- |
|  | No. | Cyan+ | Cyan- | Cyan- |
|  |  | DsRed+ | DsRed- | DsRed+ |
| C-R+ G0 Male #1 X WT Females | 246 | 0 | 123 | 123 |
| C-R+ G0 Male #2 X WT Females | 163 | 0 | 86 | 77 |
| C-R+ G0 Male #3 X WT Females | 275 | 0 | 141 | 134 |
| C-R+ G0 Male #5 X WT Females | 119 | 0 | 58 | 61 |
| C-R+ G0 Male #6 X WT Females | 339 | 0 | 170 | 169 |
| C-R+ G0 Male #9 X WT Females | 116 | 0 | 56 | 60 |

ECFP+, DsRed+ individuals, which are parental in phenotype, were retained from each injection group, separated by sex and then out-crossed *en masse* to wild-type mosquitoes to test for the presence of germline mutations in the germline ECFP coding sequence. Wild-type G_0_ were excluded from further crosses. ECFP-, DsRed+ G_1_ offspring were recovered from one male outcross from a saponin injection group and one female outcross from a saponin negative injection group, 0.82% and 0.53% of total G_1_ from each group respectively. The presence of ECFP-, DsRed+ progeny from ECFP+, DsRed+ G_0_ mosquitoes likely represent editing of the G_0_ germline without visible editing of the somatic tissues.

A panel of four primer pairs was used to characterize ECFP-, DsRed+ modifications (Fig 3A). Two primer sets amplify across the genomic DNA and the left and right piggyBac arms of the transgene and so validate the presence of the transgene at the 26.10 locus. Similarly, a PCR amplicon using a primer set across the transgene insertion site indicates the presence of a wild-type chromosome at that locus. A primer set designed to amplify 984 bp across the ECFP open reading frame and all three sgRNA target sites diagnosed the mutation character at this site: out of the 19 ECFP-knockout individuals that were molecularly characterized, only two produced diagnostic bands with ECFP primers and sequencing of the amplicons revealed small indels (Fig 3B, second panel and Fig 3C, S1). In 17 cases, PCR and sequencing further confirmed the presence of the transgene at the original locus, but no diagnostic band was seen from the ECFP open reading frame (Fig 3B, third panel, S1). In one of these, no diagnostic band was seen for amplification of either the ECFP open reading frame or the left arm, but the right arm was validated as intact, indicating a disruption of at least 1600 nucleotides of sequence (Fig3B fourth panel). These results suggest that large deletions commonly occurred using ReMOT Control, similar to results observed from embryo injection of the same components.

**Figure 3:**
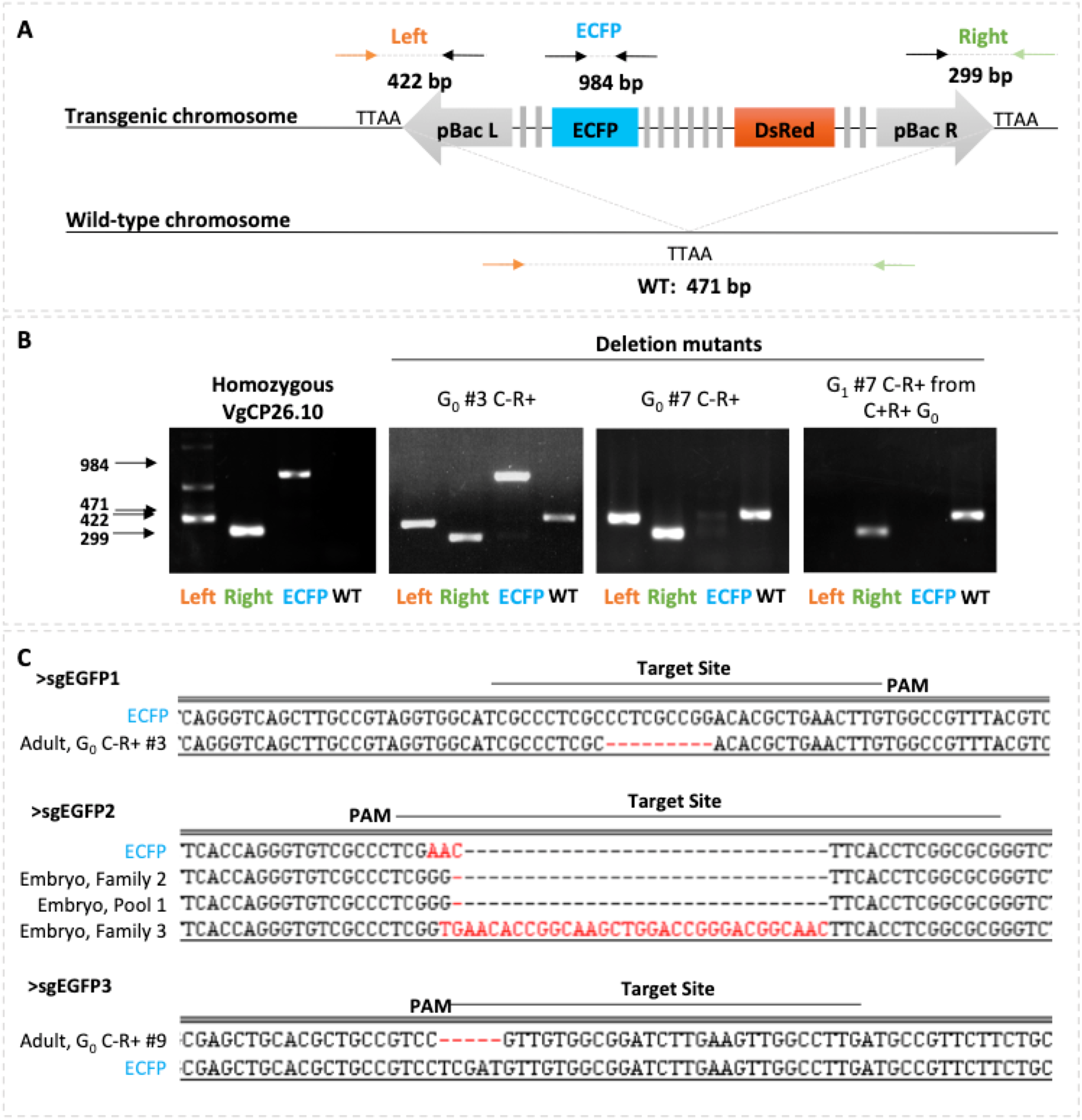
Molecular characterization of CFP negative G_0_ and G_1_ reveals both small and large deletions. A) Schematic of PCR strategy for mutant genotype characterization. Left and Right Primer sets produce amplicons of 422 and 299 base pairs, respectively, that confirm the presence of the transgene at the 26.10 genomic location. ECFP amplicon, 984 bp, spans all three target sites. Primers against genomic DNA (orange and green arrows) produce a 471bp diagnostic band from a non-transgenic chromosome. B) Representative gel images from individuals with intact transgene on both chromosomes (first panel), diagnostic amplicons for both transgenic and wild-type chromosome (second panel), amplicons diagnostic for a deletion that interrupts the primer binding site for ECFP amplification (third panel) and amplicons diagnostic for a deletion that interrupts primer sites for the ECFP amplicon and the Left amplicon (fourth panel). C) Alignments of small deletion mutants from both embryo and adult injections to ECFP. Abbreviations are the same as Figure 1 and WT: wild-type, C: cyan, R: Red, PAM: protospacer motif.

Altogether, we demonstrate with these experiments that heritable targeted knockouts can be achieved by ReMOT Control in *An. stephensi*, and that targeted editing occurs both before and after embryogenesis as evidenced both by the presence of mosaics in the G_0_ progeny and the recovery of ECFP-,DsRed+ G_1_ individuals from ECFP+,DsRed+ G_0_ outcrosses. Compared to embryo injection, ReMOT Control will allow non-specialist laboratories to conduct gene editing experiments in *Anopheles* mosquitoes and increase the number of reverse genetic studies in this malaria vector. The translation of ReMOT Control from *Ae. aegypti* to *An. stephensi* mosquitoes required modifications of the injection parameters, including timing, concentrations, and endosome escape strategies. The use of saponin dramatically increased the efficiency of knockout, consistent with results of analogous injections of the endosomal escape reagent chloroquine in the application of ReMOT Control to *Ae. aegypti*. It is possible that many reagents that mediate endosomal escape are usable for ReMOT control, which will further the adaptability of this technique. The ligand P2C is broadly effective in mosquitoes^8^, and may be useful in other insect species as well. In organisms where P2C is not effective in delivery, other ligands may be identified from proteins intrinsic to that species that are specifically targeted to the germline. We anticipate that such developments along with the straight-forward set of modifications for the adaptation to *An. stephensi* demonstrated here, will lead to the application of P2C-Cas9 to genetic studies not only to other important malaria vectors, but also to other species currently recalcitrant to gene editing techniques.

## Methods

### Mosquitoes

*Anopheles stephensi* wild type (Liston strain) and *Anopheles stephensi* transgenic line VgCp26.10 were reared at 27°C, 75±10% relative humidity, 12 h light: 12 h dark in a walk-in environmental chamber. Larvae were fed with a slurry of 1:2 by volume Tetramin:baker’s yeast. Adults were provided *ad libitum* with 10% sterilized sugar on a cotton ball. For injection experiments female mosquitoes were fed on anonymous human blood (Biospecialty Corp.) using a water-jacketed membrane feeding system.

### Embryo Microinjections

*Anopheles stephensi* females from a homozygous transgenic line VgCp26.10^7^ were used to generate heterozygous embryos for injection. Blood-fed *An. stephensi* mosquitoes were induced to lay eggs 3-5 days after a blood meal by combining 6-10 females in a narrow *Drosophila* vial with cotton and Whatman filter paper wet with isotonic buffer (150mM NaCl, 5mM KCl; 10mM HEPES; 2.5mM CaCl_2_; pH 7.2). At approximately 1 hour and 15 minutes after laying, a paintbrush was used to transfer embryos that were sufficiently melanized to a small piece of filter paper wet with isotonic buffer. Under a dissecting scope, the outer membrane was removed with jeweler’s forceps from the embryos and embryo were aligned with posterior poles to the left while maintaining moisture on the paper. After alignment of 30-70 embryos, the paper was dried briefly, embryos were transferred to toupee tape on a glass slide and an oil mixture (1:1 Halocarbon 700 oil: Halocarbon 27 oil) was used to cover the embryos to prevent further desiccation. Quartz needles were pulled using a Sutter P2000 needle puller and were used with a Femtojet injector (Eppendorf) and InjectMan micromanipulator. Following injection, oil was removed from embryos with a Kim Wipe and embryos were submerged on the tape into a petri dish with isotonic buffer and transferred back to an insectary to hatch. Hatching was monitored daily for 14 days and hatched larvae were immediately transferred to a pan with food. The injection mix from successful experiments comprised 200ng/uL *Streptococcus pyogenes* Cas9 (PacBio) and 100ng/µL each of three single guide RNAs initially described for use in EGFP targeting in human cell (12, S4). Injected embryos (generation 0, G_0_) that survived to adulthood were outcrossed: families comprised five G_0_ males allowed to mate individually to 10 wild-type females each for 2 days, then combined and pools comprised age-matched G_0_ females batch mated to wild-type males. G_1_ progeny resulting from families and pools were screened for both red and cyan fluorescence in the eyes.

### Adult Injections

Adult females with ages ranging from 5 to 22 days old were blood-fed using a glass water jacketed membrane feeder. The next day, females were immobilized by incubation at 4°C until motionless, then placed on ice and sorted. Females with visible blood-meals were injected intrathoracically using a glass needle drawn from a glass capillary (World Precision Instruments) using a needle puller (Sutter P2000) and aspirator assembly (A5177, Sigma) until we could observe visible distention of the abdomen, diuresis or liquid emerging from the injection site (approximately 200 nl).

### Adult Injection Mix Preparation

P2C-Cas9 and P2C-EGFP proteins were expressed from pET28a-P2C-Cas9 and pRSET-P2C-EGFP respectively by recombinant BL21 E.coli (NEB) as described in detail in Chaverra-Rodriguez et al.^8^. Single guide RNAs (sgRNAs) were prepared using a PCR template amplified by CRISPR_F primers designed for each ECFP target and CRISPR_R primers (S4. *In vitro* transcription was used to produce sgRNAs from PCR template using either MegaScript T7 or MegaScript RNAi kits with at least 1000ng of PCR template. Two to four volumes of the *in vitro* transcription reaction were required to achieve the high quantities of guide RNA required for adult injections. In addition to P2C-Cas9 and sgRNAs, saponin was included in some injection mixes. Saponin dilutions were prepared fresh prior to each injection from dry crude extract of *Quillaja* bark containing >20% saponin (Sigma-Aldrich). All concentrations of saponin reported here represent concentration of crude extract.

The final step in P2C-Cas9 expression and purification from transformed *Escherichia coli* is dialysis of the protein a pH 8.0 buffer consisting 50 mM Tris-HCl pH 8.0, 200 mM KCl, 0.1 mM EDTA, 0.5 mM phenylmethylsulfonyl fluoride (PMSF) and 1mM dithiothrietol (DTT). Injections using this buffer produced a high mortality in *An. stephensi* females but we found that concentrated stocks of P2C-Cas9 protein often precipitated when water, 1X PBS or guide RNA solutions were added to the protein solution. However, if the P2C-Cas9 protein was diluted in a large volume (300-500 µL) of 50 mM Tris-HCl pH 8.0, 200 mM KCl and small volumes (∼ <20 µL) of highly concentrated (∼ 3-10 µg/µL) sgRNA was added and allowed to complex, the buffer of the RNP could then be exchanged with 1X PBS using an Amicon 10K filter column, which retains the large protein and both bound and unbound guide RNA. Following complexing of the RNP and buffer exchange, saponin, water and additional sgRNA were added to produce solutions of 0.5-0.75 µg/µL P2C-Cas9, 0.75-1.2 µg/µL of total guide RNA (from a premixed solution of the three ECFPs), 50 mg/L saponin extract from *Quillaja* bark and less than 5 µM residual KCl.

### Larval Screening

G_0_ and G_1_ larvae were screened at larval instar stage 3 or 4. Larvae were immobilized by applying larvae to a wet filter paper in a Büchner funnel attached to vacuum filtering flask. Larvae were kept slightly wet during screening under UV fluorescence microscopy with a Leica dissecting scope or Zeiss Axiozoom.

### Imaging

To determine the optimal time for P2C-mediated delivery, adult females were injected with P2C-EGFP^8^ 24 hours before and 4, 24, and 48 hours after a bloodmeal. Adult female ovaries were dissected 72 hours post-blood meal, mounted on a slide in SlowFade™ Gold Antifade Mountant (Invitrogen) between two double layers of scotch tape to prevent the ovaries from being squashed by the plastic coverslip. The coverslip was sealed in place using nail polish on the edges of the slip and ovaries were visualized on Olympus BX41. All images were captured using 311ms exposure for 200X and 1030ms exposure for 40X images.

### Molecular analysis of mutant individuals

Putative mutant G_0_ and G_1_ mosquitoes were collected individually and G_1_ progeny from mutant G_0_ were collected in groups of five individuals, frozen and stored at -20°C. Genomic DNA was extracted using Wizard^®^ Genomic DNA Purification Kit or Qiagen DNeasy^®^ Blood & Tissue kits following the protocol for extraction of DNA from animal tissues. Genomic DNA was used as a template for PCR using the NEB Phusion enzyme and using primers against the transgenes and surrounding DNA (Fig 3, S4). Diagnostic gel bands were extracted using Zymoclean Gel DNA Recovery kit (Zymo, Irvine) cloned into pJet1.2/blunt using the CloneJET PCR Cloning Kit (ThermoFisher) and sequenced. Sequence analyses were performed using the DNAstar Lasergene suite.

## Acknowledgments

This work was funded by NSF/BIO grant 1645331, NIH/NIAID grants R21AI111175 and R01AI128201, USDA/NIFA grant 2014-10320, USDA Hatch funds (Accession #1010032; Project #PEN04608), and a grant with the Pennsylvania Department of Health using Tobacco Settlement Funds to JLR, USDA/NIFA grant 2017-67012-26101 to VMM, Wolfson Foundation and Royal Society fellowship RSWF\R1\180013, and NIH/NIAID grants R21AI138074 and R21AI129507 to GLH, and NIH/NIAID grant R01AI29746 to AAJ.

## Figure Legends

**Supplemental Figure 1:**
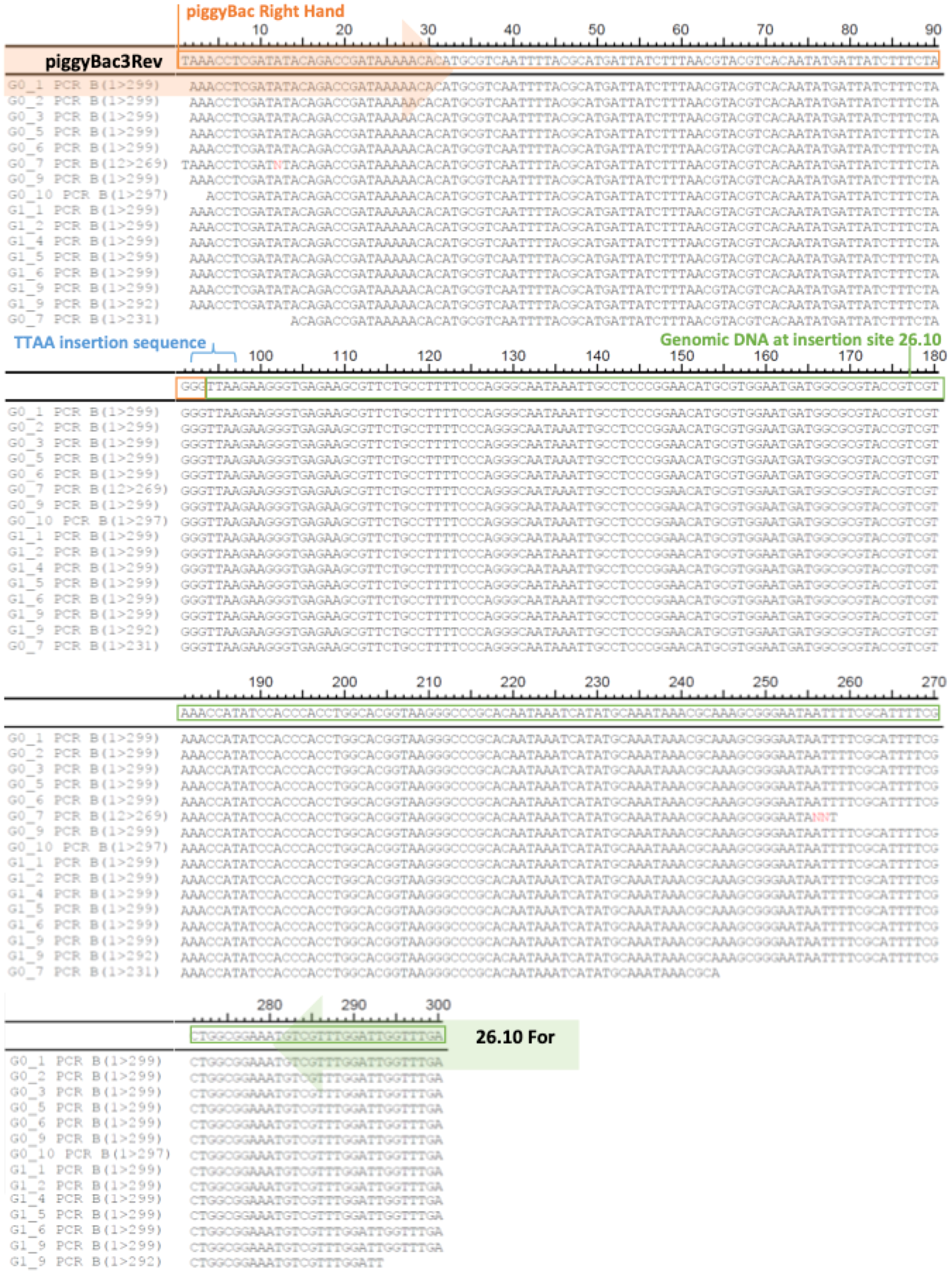
Alignment of Amplicon sequences from PCR B validate the presence of the transgene in ECFP knockout individuals

**Supplemental Table 1:**
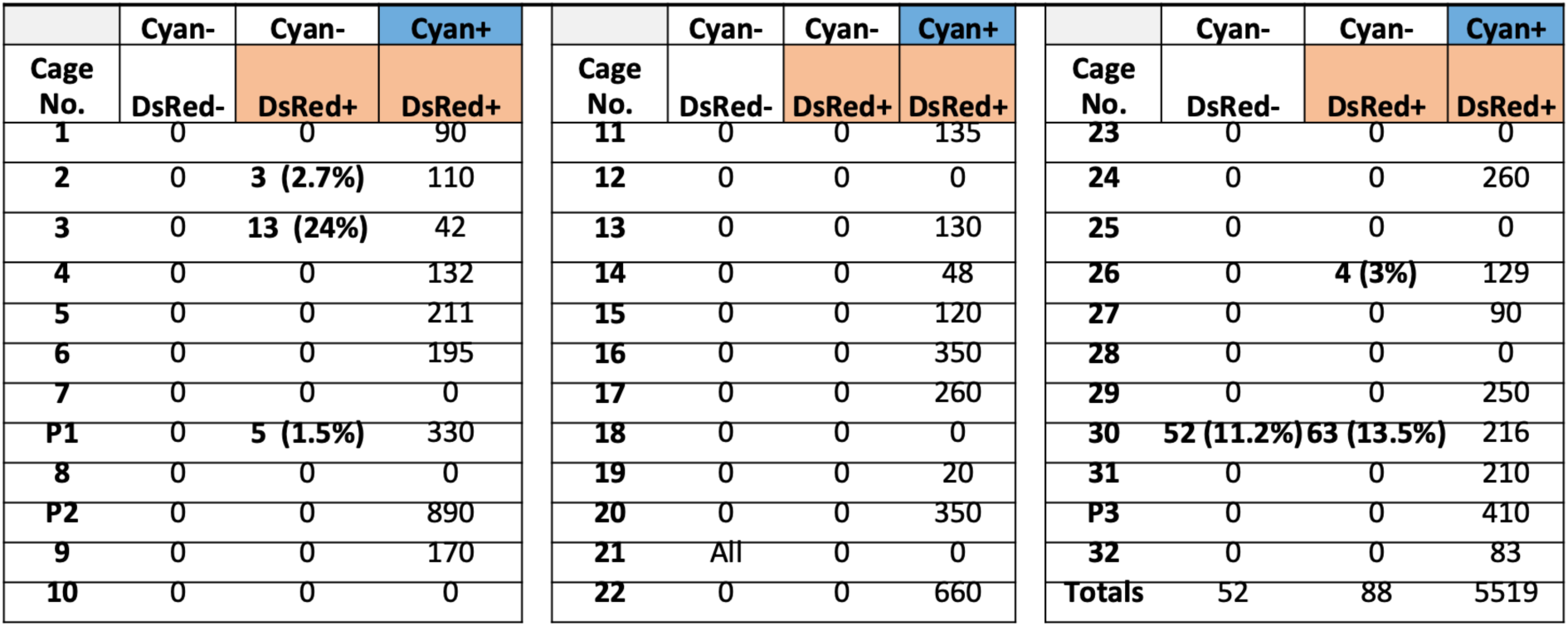
Results from screening Single G_0_ male families (1,2,3 etc) and pools of G_0_ females injected as embryos with 200 ng/μL Cas9 and three sgRNAs targeting ECFP (300ng/μL total)

|  | Cyan- | Cyan- | Cyan+ |
| --- | --- | --- | --- |
| Cage No. | DsRed- | DsRed+ | DsRed+ |
| 1 | 0 | 0 | 90 |
| 2 | 0 | 3 (2.7%) | 110 |
| 3 | 0 | 13 (24%) | 42 |
| 4 | 0 | 0 | 132 |
| 5 | 0 | 0 | 211 |
| 6 | 0 | 0 | 195 |
| 7 | 0 | 0 | 0 |
| P1 | 0 | 5 (1.5%) | 330 |
| 8 | 0 | 0 | 0 |
| P2 | 0 | 0 | 890 |
| 9 | 0 | 0 | 170 |
| 10 | 0 | 0 | 0 |
| Cage No. | DsRed- | DsRed+ | DsRed+ |
| 11 | 0 | 0 | 135 |
| 12 | 0 | 0 | 0 |
| 13 | 0 | 0 | 130 |
| 14 | 0 | 0 | 48 |
| 15 | 0 | 0 | 120 |
| 16 | 0 | 0 | 350 |
| 17 | 0 | 0 | 260 |
| 18 | 0 | 0 | 0 |
| 19 | 0 | 0 | 20 |
| 20 | 0 | 0 | 350 |
| 21 | All | 0 | 0 |
| 22 | 0 | 0 | 660 |
| Cage No. | DsRed- | DsRed+ | DsRed+ |
| 23 | 0 | 0 | 0 |
| 24 | 0 | 0 | 260 |
| 25 | 0 | 0 | 0 |
| 26 | 0 | 4 (3%) | 129 |
| 27 | 0 | 0 | 90 |
| 28 | 0 | 0 | 0 |
| 29 | 0 | 0 | 250 |
| 30 | 52 (11.2%) | 63 (13.5%) | 216 |
| 31 | 0 | 0 | 210 |
| P3 | 0 | 0 | 410 |
| 32 | 0 | 0 | 83 |
| Totals | 52 | 88 | 5519 |

**Supplemental Table 2:**
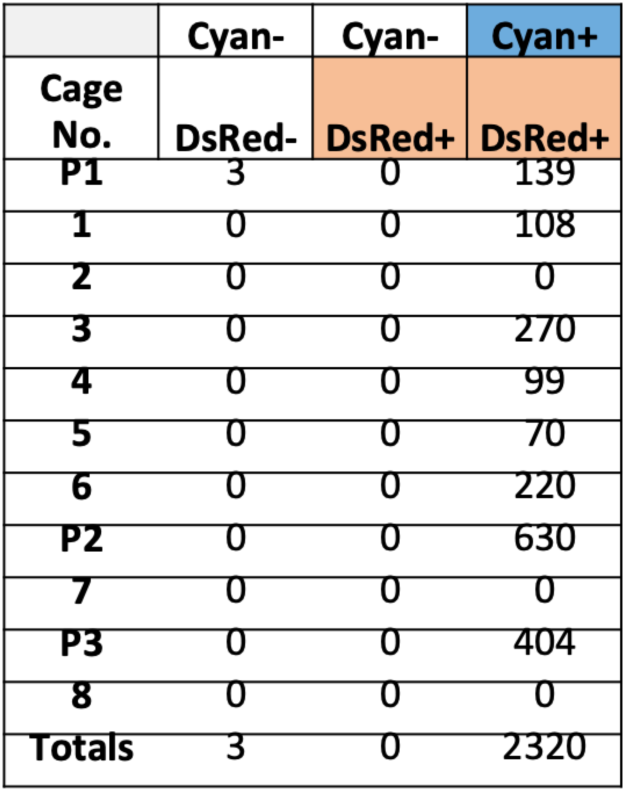
Results from screening Single G_0_ male families (1,2,3 etc) and pools (PX) of G_0_ females injected as embryos with 200 ng/μL Cas9 and three sgRNAs targeting ECFP (total 75ng/μL)

**Supplemental Table 3:**
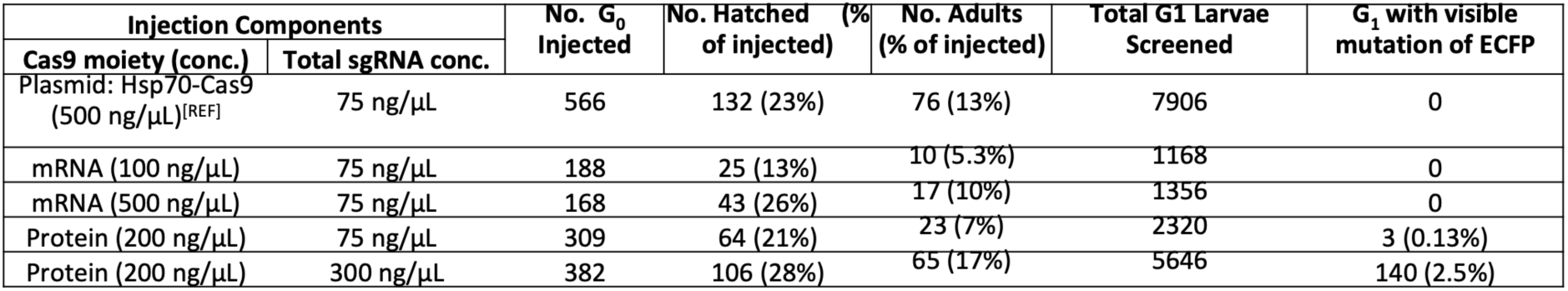
Summary of Injection components used for G_0_ embryo injections and resulting gene-edting detected in G_1_

**Supplemental Table 4:**
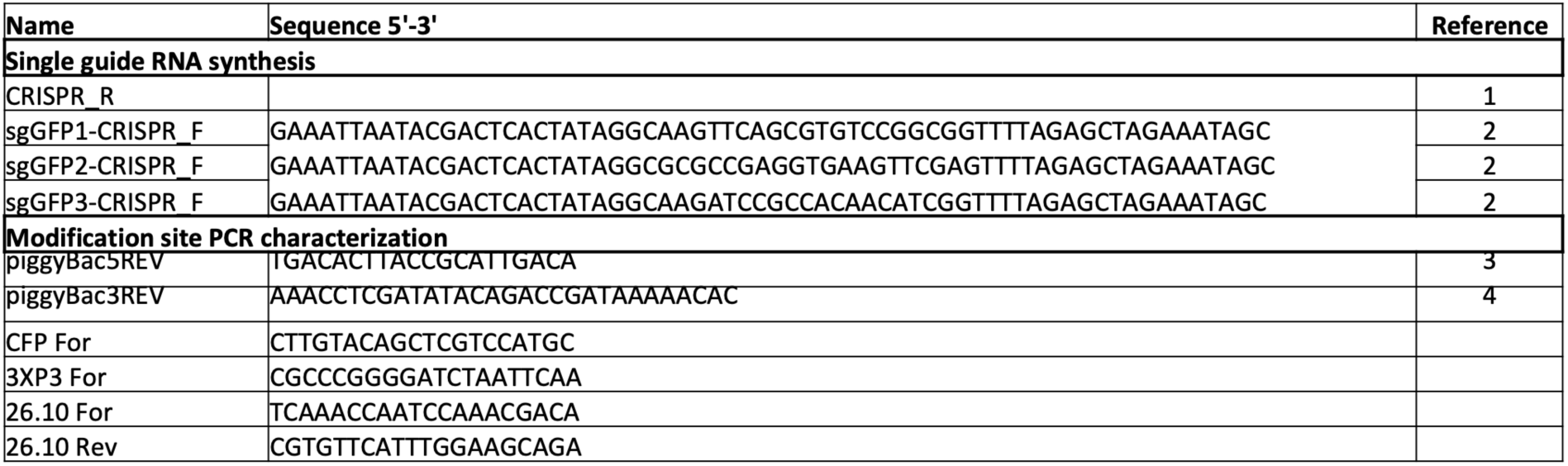
primers used for current study.

